# Robotic System for Organoid Assembly in a Multi-Well Microfluidic Chip

**DOI:** 10.1101/2023.10.02.560601

**Authors:** David M Sachs, Kevin D Costa

**Affiliations:** Cardiovascular Research Institute, Icahn School of Medicine at Mount Sinai, New York, NY; Department of Genetics and Genomic Sciences, Icahn School of Medicine at Mount Sinai, NY, NY

## Abstract

While many cell culture systems are sensitive to the conditions in which cells are introduced into the system, we find that in situ differentiated tube-shaped microfluidic organoids have a particularly high sensitivity. Preliminary experiments using conventional seeding techniques revealed that biases in initial cell number and distribution dramatically impacted organoid shape and behavior downstream. Residual flows during seeding further complicated the process, dispersing cells to undesirable locations within the chip. To address this problem, a a robotic seeding system for controlling the process of inserting cells into microfluidic chips was developed. Environmental control of temperature, CO_2_, and humidity was implemented by modifying a commercial Arduino-controlled incubator. An eight-channel syringe pump controlled flow to eight cell dispensers, while a vertical leadscrew stage raised and lowered them, and a set of stackable flexure micromanipulators individually controlled the X and Y position of each cell dispenser. The flexure manipulators were 3D printed, driven by low-cost motors and electronics, and required little assembly and no alignment, resulting in a cheap and scalable method of controlling a dense array of micromanipulators. A dual objective microscope on a motorized gantry used an oblique lighting system to observe the seeding process, allowing for real-time interventions or passive observation of automated protocols. The robotic cell seeding system provided a platform for optimizing a sensitive process towards increasing the repeatability and physiological relevance of tube-shaped microfluidic organoids.

## Introduction

In all cell culture systems, the behavior of cells is heavily dependent on the method of introducing the cells and the initial conditions present in the system. Cell seeding typically involves practical but non-physiological processes as the cells are removed from their previous environment in order to be inserted into a new environment. This may involve enzymatic digestion of matrix and membrane attachment proteins, removal of calcium from the media, exposure to room temperature or colder, pH changes due to atmospheric CO_2_ exposure, shear stress, centrifugation, trituration, and dispersal. Poorly optimized or variable cell seeding methods result in modulated behavior downstream.

With standard cell culture plates, plating density is known to be an important factor for downstream cell behavior. iPSCs have been shown to have different lineage biases that depend on initial plating density.^1^ Other environmental factors such as temperature, oxygen content, oxidative stress, and shear stress have been shown to induce a variety of transcriptional changes and differentiation of iPSCs.^2^ Passaging techniques for iPSCs often involve leaving the cells attached in small clumps rather than digesting and triturating them to single cell suspensions, leading to higher viability, but producing an additional variable of the size of the clumps.^3^ Adherence and viability can be improved via the addition of a ROCK inhibitor.^4^ In addition, cell behavior during seedings is temperature dependence, as temperature impacts both cell clumping^5^ and cell adherence to the destination surface.^6^

The seminal intestinal organoid developed by Sato et al.^7^ required extracting Lgr5+ cells from mouse crypts and plating them at one cell per well, with 6% of the cells developing into intestinal organoids. Arora et al. later increased the yield by taking advantage of the fact that, as the single cells begin to form pre-organoid multicellular aggregates, the aggregates successfully forming intestinal organoids can be identified automatically due to aggregate size and inner mass size, and harvested and replated with high viability, allowing for a second seeding with higher yield.^8^ For iPSC derived organoids, an additional selection layer is sometimes necessary, as not all pluripotent cells will develop into the desired organoid precursors. Spence et al. cultured iPSCs in a monolayer in Matrigel and differentiated them into endoderm precursors, some of which budded off into floating hindgut spheroids.^9^ In this case, the budding process serves as an enrichment step, as the floating spheroids could then be harvested and re-plated.

That these processes had such a low yield, yet produced ground-breaking successful results, highlights an advantage of organoid culture systems that use standard cell culture plates and can rely on multiple passes for enrichment. In contrast, in an in situ differentiated organoid-on-chip system with many interconnected organoids, all organoids must have very high yield in order for the chip to have reasonable yield. One way to improve the microfluidic chip yield would be to form the organoids in standard cell culture plates and then seed the organoids into the chip.^10^ This may be appropriate if spheroids are acceptable, but not if organoid tubes or other complex morphological structures are to differentiate with guidance from the mechanical constraints of the chip itself.

Drakhlis et al.^11^ generated iPSC aggregates by plating 5,000 cells in U-shaped ultra-low attachment plates and centrifuging the plates. At day 2, the aggregates were replated into Matrigel droplets, where they were differentiated into complex foregut containing heart organoids. This process had an efficiency of 80%, demonstrating that high yield organoids can be generated from pluripotent cells. The efficiency relied on an accurate iPSC seeding process, as aggregates of 3,500 cells or 10,000 cells did not form organoids.

Microfluidic cell culture systems leverage a variety of loading techniques depending on the types of cells being loaded and the desired behavior. In the seminal Huh et al.^12^ lung on chip system, human alveolar epithelial cells and human pulmonary microvascular endothelial cells were seeded into separate compartments separated by a membrane, and grown to confluence. Culturing confluent monolayers of proliferative cells allows for a less precise cell loading technique with high tolerance for variability, as the monolayer will grow into any empty space, and in fact these cell were simply loaded into syringes and injected into the chip by hand.^13^ Higher throughput chips require parallel cell seeding into many microfluidic regions. Mimetas, an organ-on-chip company, provides a chip with up to 96 culture units, in which cells are introduced into inlets, and perfused into the microfluidic region by rocking the chip.^14^ In an extreme example, the Berkeley Lights platform contains 3500 NanoPens per chip and allows cells to be maneuvered into the pens in parallel via OptoElectroPositioning, after which the cells can grow into small colonies within the pens.^15^

Some microfluidic systems seed cells at inlet ports that are separate from the final culture chamber, and these systems must address sedimentation problems that result in cells adhering before arriving the desired destination. Yun et al. remedied this by adding complexity to the chip via other fluid layers to separate the cells from surfaces before they arrived at culture chambers.^16^ Patra et al., in contrast, leveraged sedimentation to trap cells in microwells for culture,^17^ but such systems must still avoid cell adhesion on route to the microwells.

Rather than transporting cells via flow to the culture location, open microfluidic chips have microstructures that are open from above, allowing cells to be introduced by pipetting and gravity sedimentation in a manner similar to plating cells in standard culture plates, as well as allowing later access to the cells from the same opening.^18^ Open well microfluidics also allows for seeding organoids preformed in standard culture plates.^10^ While completely open microfluidic chips are limited in their ability to have gradients of signals within the media, as the open channels and wells would be covered by a shared reservoir, a partially open chip design could use open wells and enclosed channels, combining the easy access of the cells within the wells with the higher shear stress and separation of media reservoirs allowed by enclosed channels.

The cell culture enclosure can also be assembled around the cells as they are seeded, rather than seeding cells into a pre-formed structure. One version of this is the hanging drop method, in which cells or spheroids are pipetted into a culture system in a drop of liquid that hangs upside down, with the surface tension and air-liquid interface generating a curved well-like culture system that can be integrated into a more complex multi-tissue array.^19^ Bioprinting is a more intricate example of this, in which cells are added to a complex structure as the structure is being generated around them. Bioprinting is an exciting and rapidly developing technology that has uses for generating large, complex, tissue constructs.^20^ It can also be used as part of the formation of organ-on-chip systems; however, the current low resolution of bioprinters (hundreds of microns) is not appropriate for the fabrication of microchannels, with the possible exception of the UV printing of 25 μm features in acellular hydrogels using digital micromirror devices.^21, 22^ This technology will be revisited as it continues to mature. In an example of bioprinting applied to organoid seeding, Lawlor et al. used a bioprinter system to extrude cells in matrix in precise spheroids, creating more reproducible kidney organoids.^23^

Introducing cells into a microdevice is a complex task that requires control of environmental and mechanical parameters, an understanding of cell behavior as a consequence of seeding perturbations, the required final cell location and density, interactions between the cells, reagents, and extracellular matrix, and the synergistic design of both the chip and the seeding process to ensure an effective delivery of the cells to their destinations. For complex in vitro systems, automation may need to be applied in order to achieve the appropriate level of control and repeatability.

## Methods

### Machine Design and Construction

Machines were modelled in Fusion 360 (Autodesk). Off-the-shelf parts were acquired from McMaster-Carr or Misumi, with custom parts 3D printed in ABS on an Ultimaker S3 3D printer, or laser cut in acrylic on an Epilog Helix. Low level control of sensors, actuators, and LEDs was performed on Arduino Neros, with separate Arduinos controlling the valves, incubator, and motors. Gantry and cell dispenser NEMA-17 (Digi-Key) stepper motors were controlled by ST L6470 motor drivers on X-NUCLEO-IHM02A1 (Digi-key) boards, with the boards configured to be stacked in a daisy chained manner. Micromanipulator and focus 28BYJ-48 motors were controlled by Darlington transistors (Digi-Key, ULN2803A), interfaced by a series of daisy chained shift registers (Digi-key, SN74HC595N). This allowed all 4 NEMA-17 motors and all 18 28BYJ-48 motors to be controlled by two SPI commands.

### Software Design and Architecture

Arduino firmware was used to perform low level interfacing to actuators, sensors, and LEDs, while accepting serial commands and outputting a stream of motor location and sensor data. Arduinos were connected to the USB ports of a Raspberry Pi, which ran a Python user interface via the Pygame library, allowing a user to control the X and Y position of the microscope, the focus, a set of servo controlled valves, the incubator, the vertical position of the cell dispensers, and the syringe pump with simple g-code based text commands and mouse movements, while visualizing the seeding process with the Raspberry Pi camera-based microscope. The text commands could be entered in real time to test adjustments to the seeding process while watching cells respond, or the commands could be assembled into scripts executed by the Raspberry Pi in order to automate complex tasks while minimizing variability and user mistakes.

## Results

Early seeding attempts were performed manually, by injecting cells into chips via a syringe or syringe pump (**Figure 1A**), positioning cells in pipette tips (**Figure 1B**) or transfer pipettes (**Figure 1C**) to sediment into the chip via gravity. Despite early successes in repeating more conventional microfluidic scenarios, such as endothelializing channels and culturing aggregates of beating cardiomyocytes on chips, novel difficulties arose when attempting to create tube-shaped organoids. In these cases, the seeded chips were mostly unusable, with cells either in the wrong parts of the chip or had somehow never reached the chip at all. When performing test seedings while observing the chips under a microscope (**Figure 1D**), it was revealed that the final location of the cells was exquisitely sensitive to movement of the chip or any interconnecting tubes or pipette tips, with even small movements or adjustments leading to forces that directed cells to the wrong parts of the chip. In addition, cell clumping during seeding, a particularly acute problem with iPSCs, could lead to cells blocking the pipette tips and never arriving at the chip at all.

**Figure 1.**
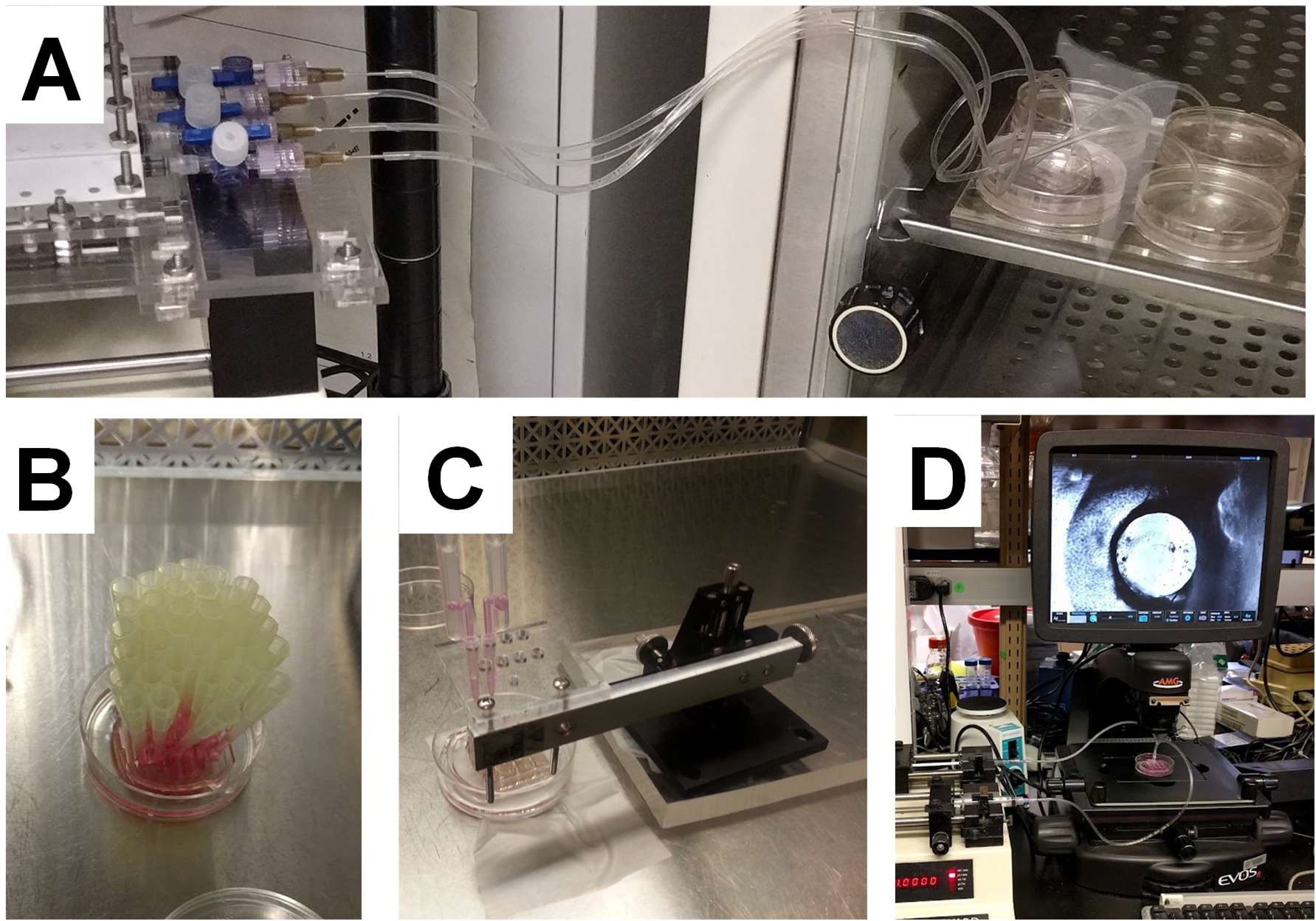
Early Seeding Attempts. **A)** HUVECs were seeded into microfluidic chips with a syringe pump (Harvard Apparatus). **B)** Cells were withdrawn into pipettes and manually embedded in the chip, allowing the cells to sediment. **C)** Cells were withdrawn into transfer pipettes and sedimented into the chip after positioning with a micromanipulator. **D)** Non-sterile seedings tested while viewing with a microscope (Evos FL).

Importantly, these struggles were a function of the project goal, which was to generate in situ differentiated tube-shaped organoids that connected to microfluidic channels as they developed, a goal that turned out to be surprisingly difficult when compared with more common goals of seeding channels with endothelial cells or generating sphere-shaped organoids. Seeding cells such that they bridge the space between two microfluidic channels, create structures connecting the channels rather than detaching from one or both of them, are free to form channel bridging structures but are not forcibly confined to do so in a way that would inhibit future morphogenesis, and to do all this with any hope of repeatable behavior, turned out to be very difficult, and success was dependent on enforcing strict control over initial cell positions. Any early bias in the initial distribution of the cells would result in detachment from one or both channels. Addressing these problems would require the construction of a more controllable seeding system.

### Environment Control During Seeding

Moving cells from a standard plate into a microfluidic chip could result in cell death or transcriptional changes that could impact downstream organoid formation. For example, iPSCs are known to be sensitive to changes in temperature, oxidative stress, and shear stress, responding with transcriptional changes and differentiation.^1–4^ Temperature control also impacts cell adherence^6^ and clumping^5^ during seedings, which is important for directing cells to specific compartments of microfluidic systems. Temperature additionally impacts hydrogel behavior, which must remain liquid during the seeding process. Gelatin-based hydrogels, such as gelatin methacrylate, must be kept warm to prevent early gelation, while collagen-based hydrogels, such as Collagen I and Matrigel, must be kept cool to prevent early gelation. For a prototyping system capable of supporting a variety of seeding methods of different types of cells and different hydrogels, controlling the environment during seeding is an important factor for controlling cell location and downstream reproducibility of organoid development.

To conduct a flexible range of seeding processes without contamination, the seeding mechanisms had to be environmentally controlled and located in a laminar flood hood. While commercial miniature CO_2_ incubators exist that could fit in a hood, such as the MyTemp Mini CO_2_ Digital Incubator (Benchmark Scientific), these are not structurally modifiable, and therefore not appropriate for a prototyping system that might need to expand in size and be controlled in unconventional ways. Recently, several labs have constructed DIY CO2 incubators,^24–28^ and the Pelling Lab implementation led to the company Incuvers that produces modifiable Arduino-controlled incubators,^29^ from which we acquired the Incuvers Model 1 (**Figure 2A**).

**Figure 2.**
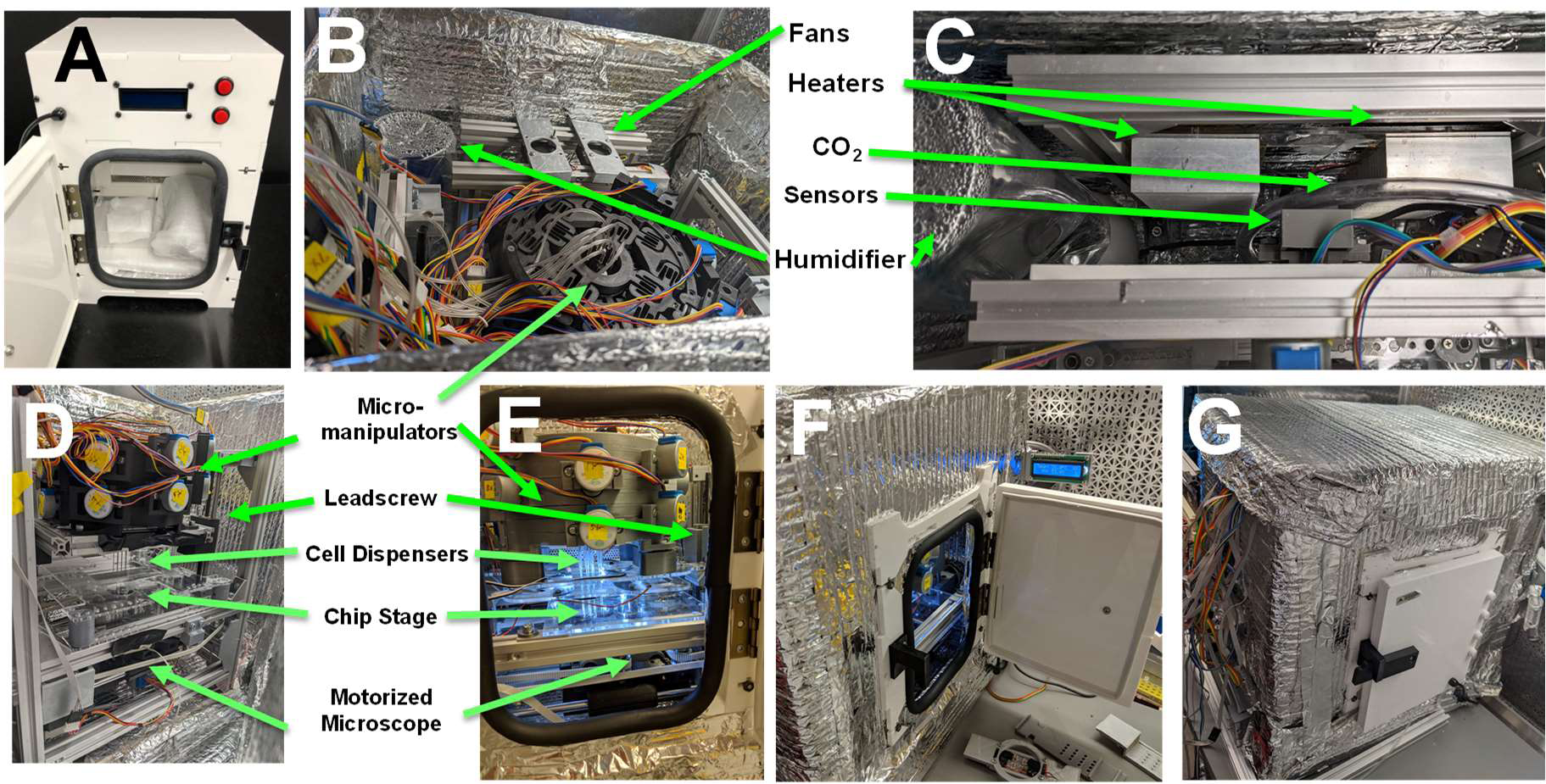
Incubated Cell Seeding Machine. **A)** The Incuvers Model 1 Incubator. **B)** Top removed for maintenance, view of the machine revealing the environmental control system (back, covered by fans) and micromanipulators (front). **C)** Top view close up of the environmental control system with the fans removed, showing heating elements, CO_2_ injecting tube, temperature/humidity/CO_2_ sensors, and humidifier. **D)** Front removed for maintenance, showing the micromanipulators, cell dispensers, one of the two leadscrews used to raise and lower the cell dispensers, chip stage, and motorized microscope. **E-F)** Assembled machine located in a laminar flow hood with the incubator door open and **G)** closed.

In order to accommodate the seeding mechanisms, the incubator had to be modified by enlarging it and adding additional entry points for maintenance. A larger frame was constructed out of laser cut acrylic sheets attached with 3D printed corner pieces. For insulation, the frame was covered in insulating aluminum foil bubble wrap (McMaster-Carr, 9367K33) and taped together with aluminum foil tape (McMaster-Carr, 7616A21). The door of the Incuvers, which contains a heating element and a temperature sensor, was repurposed as the door to the seeding machine. The top of the seeding machine and front wall were designed to be removable to allow easy access to the seeding mechanisms for maintenance from the top (**Figure 2B-C**) or the front (**Figure 2D**). The final assembly housed the seeding mechanisms and fit in a laminar flow hood (**Figure 2E-G**). This allowed the microfluidic chip to be prepared in the sterile hood, quickly inserted into the seeding machine where the cells were seeded and adhered in an incubated environment. At the end of the seeding process, the chip was removed back into the sterile environment where any final reagents were added before covering it and transporting it to the final standard CO_2_ incubator.

The provided source code for the Arduino based Incuvers control system regulated heat via heating elements and temperature sensors, and CO_2_ from a standard CO_2_ tank via a solenoid valve and a CO_2_ sensor (**Figure 2B-C**).^29^ However, humidity was controlled passively with a water pan. Because of the need to rapidly humidify and dry out the seeding machine before and after seedings, this was replaced by a controlled humidity system using a miniature ultrasonic humidifier (Fancii personal humidifier) (**Figure 2B-C**) connected to a relay-controlled outlet (Adafruit 2935) and a humidity sensor (Honeywell HIH8120-021-001). The Incuvers firmware had only rudimentary serial data output for debugging, which was modified to provide full input and output control via the USB serial port, allowing a single Raspberry Pi system to control all aspects of environmental control in addition to the other features of the cell seeding system.

After modifying the Incuvers incubator, the system had to be verified to ensure that environmental parameters were still being controlled successfully. Our modified Incuvers had significant added control challenges compared with the commercially provided version, including a bigger internal space, a custom frame that had wires and tubes passing through spaces in the walls, and the flow of air passing around it from the laminar flow hood. As the final machine was unable to maintain environmental control while the hood was on, the hood was switched off immediately after inserting the microfluidic chip. With the hood off, temperature at the location of the chip was calibrated to 37 °C using a NIST traceable temperature sensor (Fisherbrand Digital Thermometer). CO_2_ was calibrated to 5% by calibrating the CO_2_ sensor (ExplorIR-WV 20% CO_2_ Sensor) to a recently professionally calibrated standard incubator. Humidity was not calibrated, but was set to an approximate value of 90%, or as high as possible without creating condensation. Environmental control was monitored over a 9 hour time period (**Figure 3**). During this time, temperature was controlled to a mean of 36.94 °C, standard deviation of 0.06 °C, and coefficient of variation of 0.15% (**Figure 3A**). CO_2_ was controlled to a mean of 5.07%, standard deviation of 0.07%, and coefficient of variation of 1.30% (**Figure 3B**). Humidity was controlled to a mean of 90.88%, standard deviation of 1.04%, and coefficient of variation of 1.14% (**Figure 3C**).

**Figure 3.**
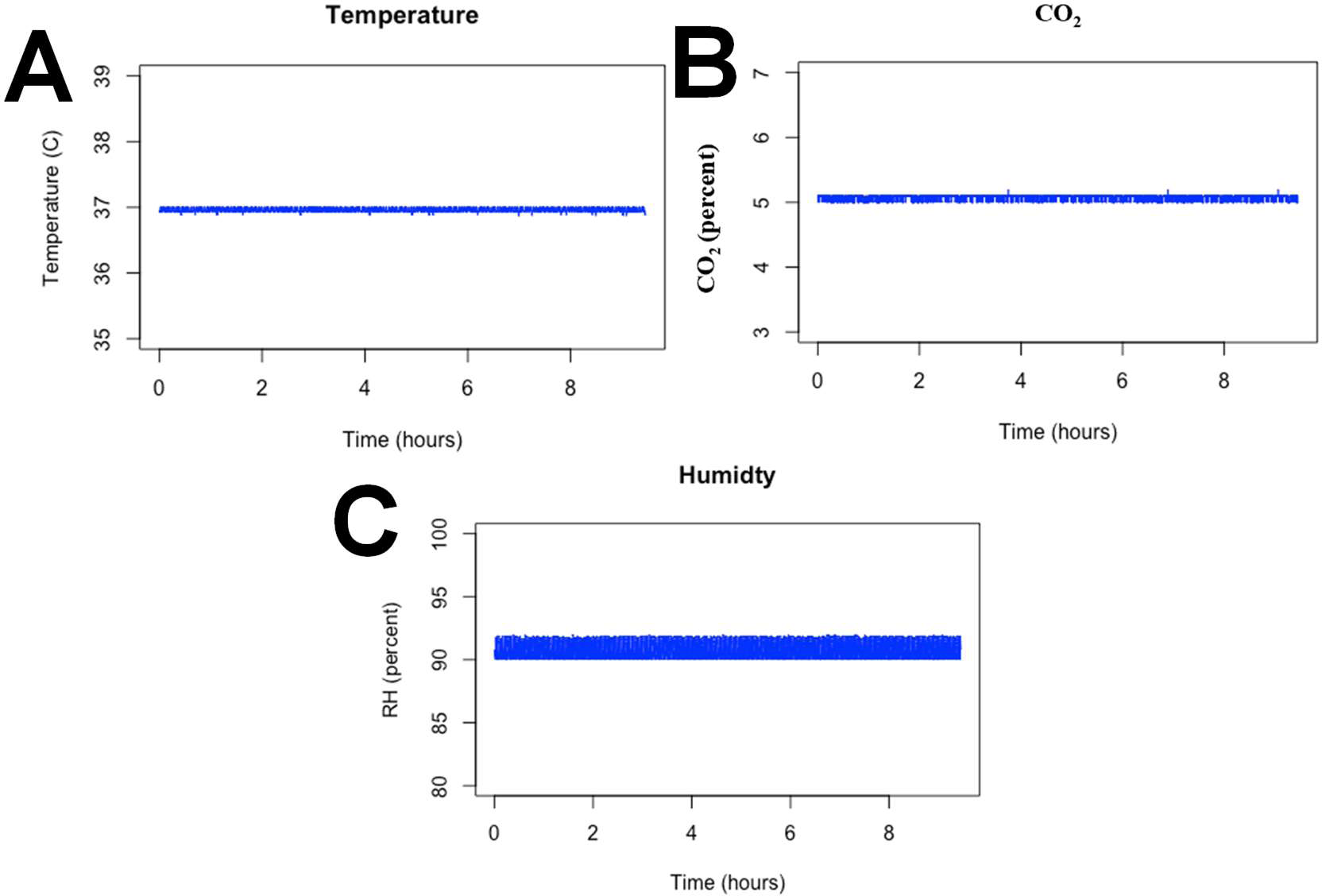
Environmental Control Stability of Cell Seeding Machine. During a 9 hour test, environmental control was measured and evaluated. **A)** Temperature was controlled to a mean of 36.94 °C, standard deviation of 0.06 °C, and coefficient of variation of 0.15% **B**). CO_2_ was controlled to a mean of 5.07%, standard deviation of 0.07%, and coefficient of variation of 1.30%. **C)** Humidity was controlled to a mean of 90.88%, standard deviation of 1.04%, and coefficient of variation of 1.14%.

While a warm environmental control was sufficient for most seedings tested, a cold environment was also evaluated, as this would enable collagen-based hydrogels that would otherwise rapidly gel before cells could be positioned. A modification to passively cool the machine was implemented by building a cooling lid containing a metal pan that held bags of propylene glycol pre-frozen in a −80 °C freezer. As propylene glycol has a freezing temperature of −59 °C, this would allow a phase change to maintain a very cold temperature. The Incuvers fans were repositioned to blow air at the bottom of the cooling pans, rapidly cooling the internal environment.

Despite the very low temperature of the propylene glycol, this process only maintained an 8 °C temperature within the seeding area, and this system was unable to prevent early gelation of collagen. If future seedings require collagen, additional experimentation would be needed to maintain the low temperature. As both air and the microfluidic chip materials are poor heat conductors, preventing collagen from gelling during seedings may require cooling control via more efficient heat conduction, by embedding the chip in a cooled aluminum block, for example. Currently, collagen is avoided as a component of extracellular matrix materials used during seedings.

### Oblique Microscope For Real-Time Seeding Feedback

When manual seedings were first attempted, they had high variability and failure rates. As it became apparent that visual feedback would be necessary for troubleshooting the seeding failures, a custom microscope was built into the seeding machine, with the initial optical design derived from the OpenFlexure microscope.^30^ The optical path of the seeding microscope consisted of a standard microscope objective, a 50 mm focal length achromatic doublet tube lens (Thorlabs AC254-050-A), and a Raspberry Pi camera (Digi-Key), using an inverted configuration for observing the chip from below. A 4X and 10X objective were mounted on a 3D printed piece that could be moved via a small screw mechanism driven by a 28BYJ-48 stepper motor (**Figure 4A**). The 4X objective was the primary workhorse for visualizing seedings, while the 10X objective was occasionally used to visualize adherent cells.

**Figure 4.**
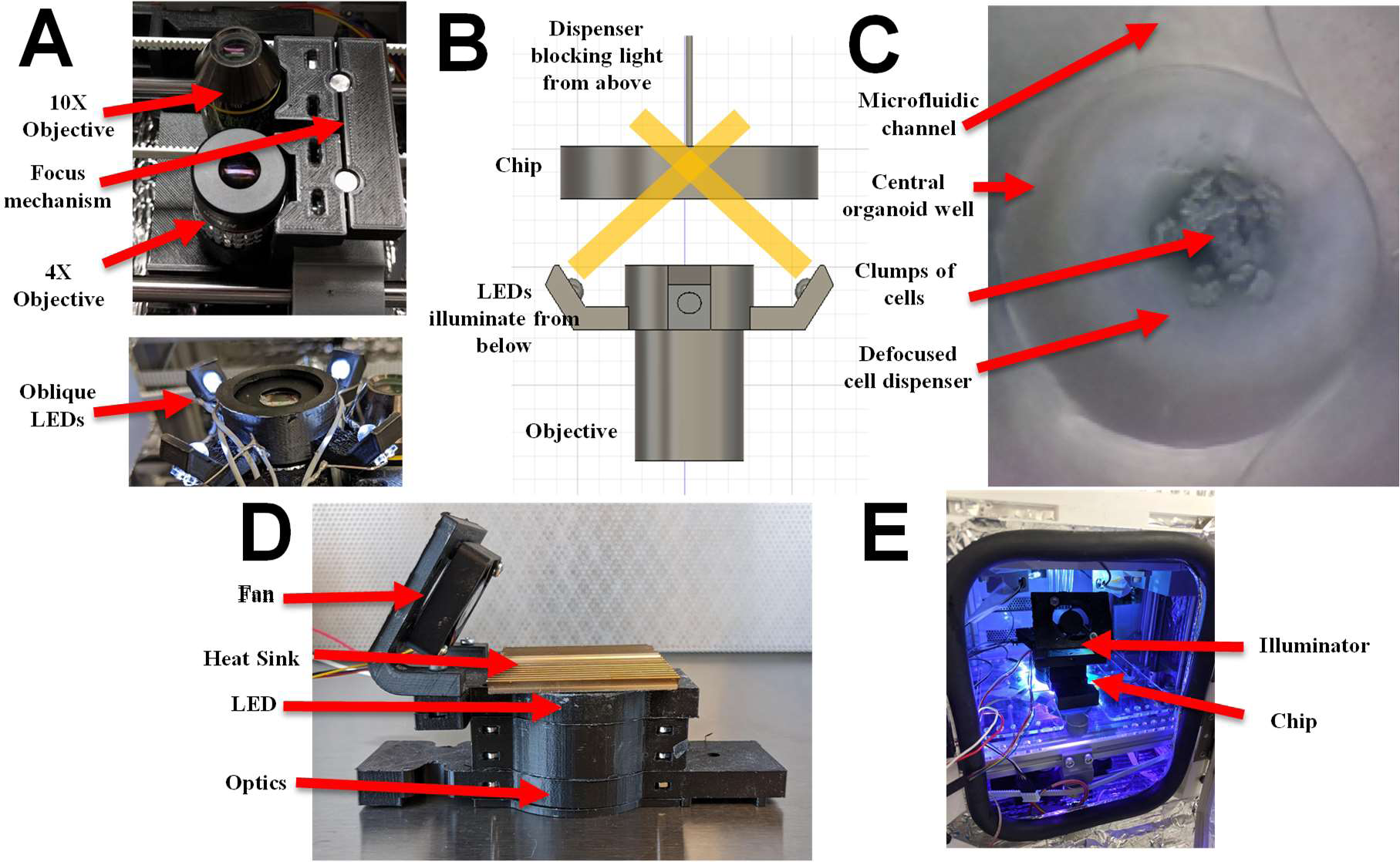
Microscope and Lighting Systems. **A)** Top: 4X and 10X objectives with focus mechanism. Bottom: ring of LEDs positioned for oblique illumination from below. **B)** Oblique lighting system diagram. **C)** Sample oblique image, showing the chip, clumps of cells, and the cell dispenser (out of focus due to being in a different focal plane). **D)** Post-seeding illuminator for bright field imaging or photoinitiation. **E)** The illuminator inserted into the machine, photoinitiating.

Because the cell dispensers blocked light from above, the seeding process was illuminated from below, with four oblique LEDs that were positioned in a ring around the 4X objective (**Figure 4A-B**). This was sufficient for visualizing clumps of cells dropping into the chip from the dispensers, as well as the dispenser locations relative to the chip (**Figure 4C**), but it was ineffective for visualizing adherent cells such as HUVECs, which are poorly visible under oblique lighting. To visualize adherent cells, the dispensers were removed after the seeding, and an additional bright field illuminator was placed above the chip. The illuminator consisted of a white LED (Mouser 749-R20WHT-F-0160), a Fresnel lens (Thorlabs FRP232), and a diffuser (Thorlabs DG20-1500) (**Figure 4D**).

For experiments involving photoinitiated hydrogels, the bright field illuminator was repurposed as a photoinitiator that could be inserted into the seeding machine after cells had been positioned in the microfluidic chip (**Figure 4D-E**). The photoinitiator contained a 10 W 523 nm LED (LED Engin LZ4-40G108-0000) for eosin Y based systems or a 40 W 390 nm LED (LED Engin LZC-C0UB00-00U4) for lithium phenyl-2,4,6-trimethylbenzoylphosphinate based systems, and an additional fan and heat sink to handle the heat generated by high power LEDs (**Figure 4D**). When using the illuminator as a photoinitiator, the diffuser was removed to increase light intensity, and when using the 390 nm LED, the plastic Frensel lens was replaced with a glass aspheric condenser lens (Thorlabs ACL50832U-A), which transmits more light at short wavelengths.

The microscope was mounted on a custom designed gantry to allow it to be moved in the x-y plane during the seeding process. The gantry had an H-bot style configuration with a belt drive controlled by NEMA 17 motors (**Figure 5**). The H-bot configuration was chosen due to its vertical compactness, with the motors mounted to the frame driving a low-profile gantry in a coupled parallel manner, and all pulleys and belts located in a single vertical plane. H-bot gantries are known to suffer from racking problems, in which high accelerations apply a torque to the frame and result in brief positioning inaccuracies;^31^ however, for this microscope application, such high accelerations are not necessary. The orthogonal focus mechanism consisted of an M3 screw driven by a 28BYJ-48 stepper motor (**Figure 5**).

**Figure 5.**
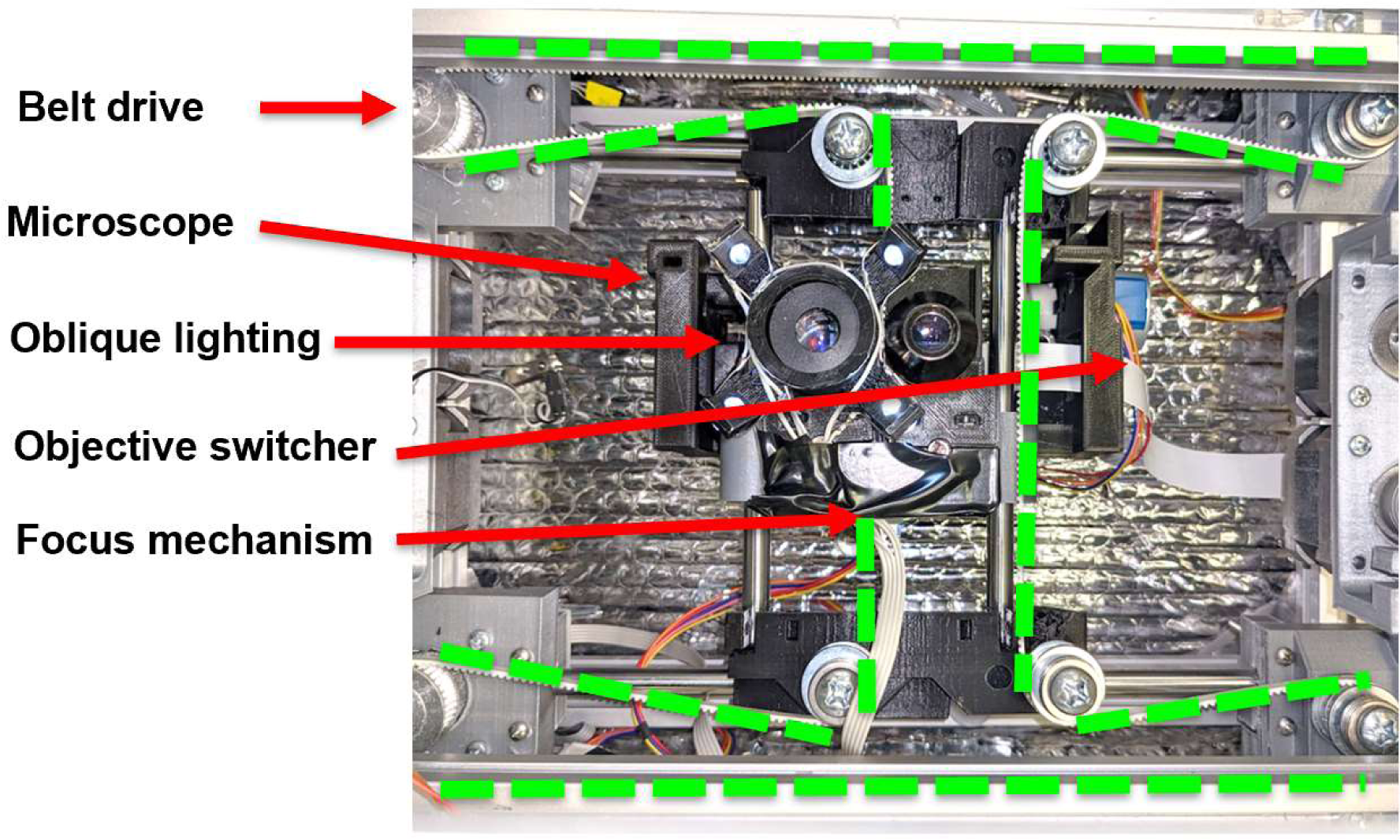
Motorized XY Gantry. H-bot belt routing (dotted lines) holding dual objective microscope with objective switcher, focus mechanism, and oblique lighting.

### Fluidic System

As the microfluidic chip was designed to have eight organoid wells that would be seeded simultaneously, an eight-channel syringe pump (New Era NE-1800) was used to withdraw cells into dispensers and deposit them into the chip (**Figure 6A**). The syringe pump was loaded with 1 mL syringes connected to tygon tubing (McMaster-Carr 6546T23, ID 1/32”, OD 3/32”) that was primed with water to minimize the air gap between the syringe pump and the cells. The tubing entered the incubated seeding machine and connected to 304 stainless steel tubes (McMaster-Carr 5560K49, OD 0.042”, ID 0.035”) with gel loading tips or additional 316 stainless steel tubes attached, described later, which were used to withdraw cells and dispense them into the microfluidic chip. The dispensers were raised and lowered with two M8 leadscrews (Misumi MTSRA8-160-S10-Q6) driven by NEMA 17 motors (Digi-Key) (**Figure 2D-E**).

**Figure 6.**
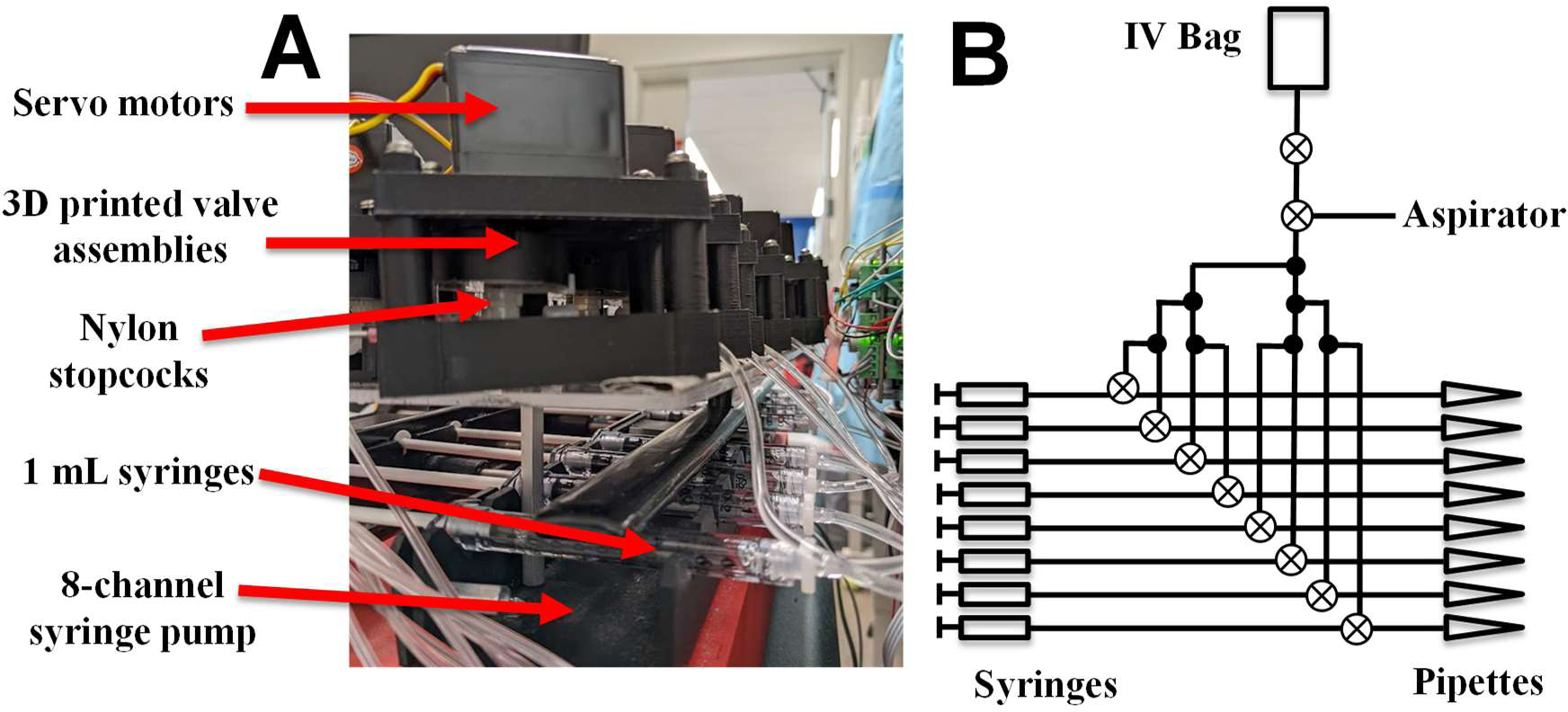
Fluidic System for Cell Dispensing and Self Cleaning. **A)** 3D printed servo-controlled valves and 8 channel syringe pump. **B)** Fluidic circuit diagram.

The seeding as well as the post-seeding cleaning process were controlled by a set of servo-controlled 3D printed stopcock valves (**Figure 6A**). As the seeding process resulted in the dispensers being exposed to cells and extracellular matrix, considerable fouling occurred that could interfere with future seedings. After seedings, cell dispensers were washed with an enzymatic detergent (Alconox, Tergazyme) followed by chemical sterilization with 2.5% trisodium phosphate with a pH of 12 (Sigma), and then distilled water. Washes were partially automated, with dispensers lowering into a 35 mm dish filled with the wash fluid, withdrawing fluid, and then repeatedly withdrawing and dispensing to apply shear stress to the contaminants. Nylon stopcocks (DWK Life Sciences, 4201634503) were used, as the more common polycarbonate stopcocks cracked with even brief contact with trisodium phosphate. Valves were controlled to sequentially drip water through each dispenser, followed by vacuum drying by sequentially connecting each dispenser to an aspirator (**Figure 6B**). The water dripping process also reprimed the syringes, allowing for a minimized and standardized air gap between the priming water and the cell laden media during the next seeding.

### Varying Seeding Parameters

Seeding parameters include the dispenser diameter and volume, the vertical movement of the dispensers with respect to the chip, and the controls sent to the syringe pump. They have a complex dependence on the cell clumping behavior, dispenser material, compliance of the fluidic system, and architecture of the microfluidic chip, and as such were designed empirically as seedings were performed with visual feedback from the gantry microscope.

Dispensers were constructed out of 316 stainless steel hypodermic tubing or polypropylene gel loading pipette tips. The gel loading tips had a 500 μm diameter tip with a 30 mm height, and a conical region that increased the volume that could be held, and minimized the height of the sedimenting cells. This resulted in 70 μl of cell laden media with a maximum height of 50 mm (**Figure 7A**). In comparison, the stainless steel tubing had a cylindrical internal volume (**Figure 7B**), with a maximum cell height of 150 mm, and a smaller volume determined by the inner diameter of the tubes. While gravity sedimentation was sufficient for seeding from the gel loading tips, the increased height and decreased volume of the stainless steel tubing required flow to push out a sufficient number of cells. In addition, the stainless steel tubing was hydrophilic, leading to strong capillary forces that pulled cell laden media up in the tube, which resulted in an air gap between the cells and the chip, preventing cells from entering. This led to a more complex seeding protocol in which a small amount of media was dispensed just prior to seeding, in order to ensure there was no air blocking the path of the cells during the seeding process.

**Figure 7.**
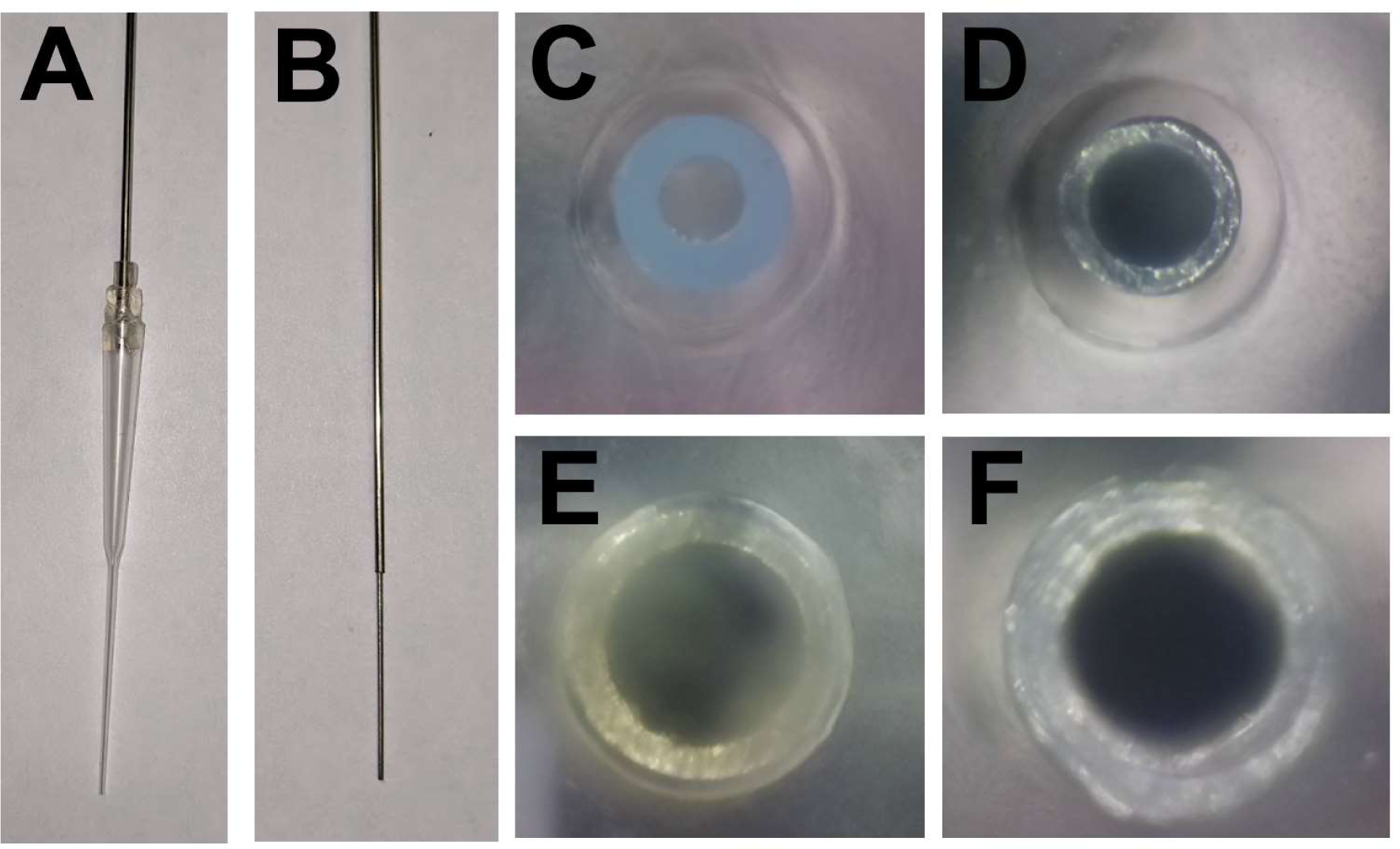
Cell Dispensers. **A)** Gel loading pipette tip attached to stainless steel tubing. **B)** All stainless steel dispenser. **C)** Gel loading tip in organoid well. **D)** OD 568 μm, ID 406 μm stainless steel tubing. **E)** OD 762 μm, ID 610 μm stainless steel tubing. **F)** OD 1067 μm, ID 686 μm stainless steel tubing.

Nevertheless, the stainless steel tubes had a number of technical advantages compared to the gel loading pipette tips. The gel loading tips have a complex assembly process, involving gluing them to a Tygon/silicone gasket attached to stainless steel tubing which could be inserted into the micromanipulator stage for dispenser manipulation; the final seal was prone to failure if any external mechanical or fluidic forces were applied to the dispenser. While the conical region of the gel loading tips could hold more media than the steel tubes, it was also prone to contamination as it acted as a trap for dust or clumps of cells, and it was difficult to clean without damaging the seal between the gel loading tip and the steel tubing. In contrast, the stainless steel tubing could be easily cleaned mechanically by reaming it with a steel wire, or chemically by pumping cleaning fluid through it. While the gel loading tip can often seed a higher density of cells using only gravity sedimentation, in practice the region between the cone and the tip is prone to cell clumping and blocking sedimentation. Finally, the cone region of the gel loading tips limits the scalability of those tips, as it limits how close together the tips can be positioned. Later versions of this system would ideally have higher throughput and would require a higher density of dispensers. Straight stainless steel tubing would be advantageous as a scalable form factor, and material properties could be modulated with PTFE or parylene coating.

Chip seeding was complicated by the presence of unpredictable residual flow, which might be generated by differential evaporation, air being released from the chip, volume changes due to extracellular matrix gelation, or other subtle changes (**Figure 8A**). Chip architectures were tested with and without an open shunt, with dramatic differences in seeding outcomes. Without the shunt, all flow within the central reservoir is directed into the organoid well (**Figure 3.8B**). This simplifies the process of directing the cells into the organoid well, and does not require that the cell dispenser be exactly positioned. However, this configuration leads to spontaneous residual flows in the organoid region, as small pressure differences in the reservoirs are directly applied to the organoid. Even after allowing the chip to settle overnight, flow was observed in the organoid region that could disturb the developing organoid. Flows generated by small displacements from the cell dispenser or pipettes could generate enough flow to disperse cells.

**Figure 8.**
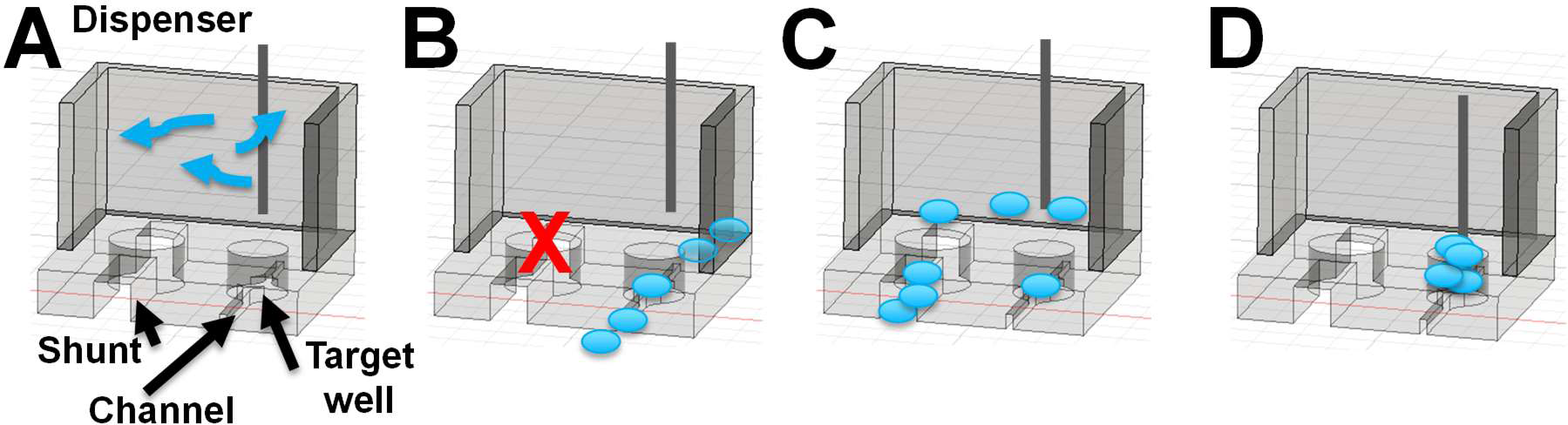
Seeding Dependence on Chip Design and Dispenser Location. **A)** Residual flows complicate seedings. B) If the shunt is closed, cells enter the channels, but are hard to control. C) If the shunt is open, and the dispenser is elevated, cells are dispersed towards or away from the shunt. D) If the shunt is open and the dispenser is in the central organoid well, cells sediment straight down into the well.

To eliminate residual flows in the chip that disturb seedings, a shunt was added that redirects any pressure difference between the central and outer reservoirs through the shunt. This allowed the organoids to be seeded and develop without interference from residual flow, but it complicated the micromanipulator positioning process, as the cells had a tendency to be directed into the shunt rather than the organoid well (**Figure 8C**). To remedy this, the cell dispenser tip must be lower in the organoid well while seeding, to shield the cell from any shunt flow (**Figure 8D**). This meant that dispensers with outer diameters larger than the central organoid well could not be used (**Figure 7F**). It also required that the cell dispenser height be accurate to within a fraction of the organoid well height (2 mm), and that the micromanipulators be capable of positioning the dispensers accurately relative to the walls of the organoid well. These requirements reinforced the need for a robotic seeding system.

The seeding process began with the preparation of a chip by coating it with extracellular matrix such as Laminin-Entactin (Corning) and filling it with the seeding media. The chip was placed in a 3D printed holder with silicone feet, to allow it to be aligned but prevent any shifting during the seeding process (**Figure 9A**). The chip holder with the chip was then inserted into the seeding machine and aligned to the dispensers using the microscope (**Figure 9B**). Cells were retrieved from six-well plates and pipetted into eight PCR tubes held by a 3D printed tube holder (**Figure 9C**) which was then inserted into the machine (**Figure 9D**). The dispensers descended into the tubes and performed a final synchronized resuspension by withdrawing, dispensing, and withdrawing again (**Figure 9E**). A final dispense of 5 μL was required to prime the fluidic system towards dispensing; without this a small air gap will form in the dispenser that blocks cells from entering the chip. The dispensers were raised and the tube holder removed (**Figure 9F**), after which the dispensers descended into the chip (**Figure 9G**).

**Figure 9.**
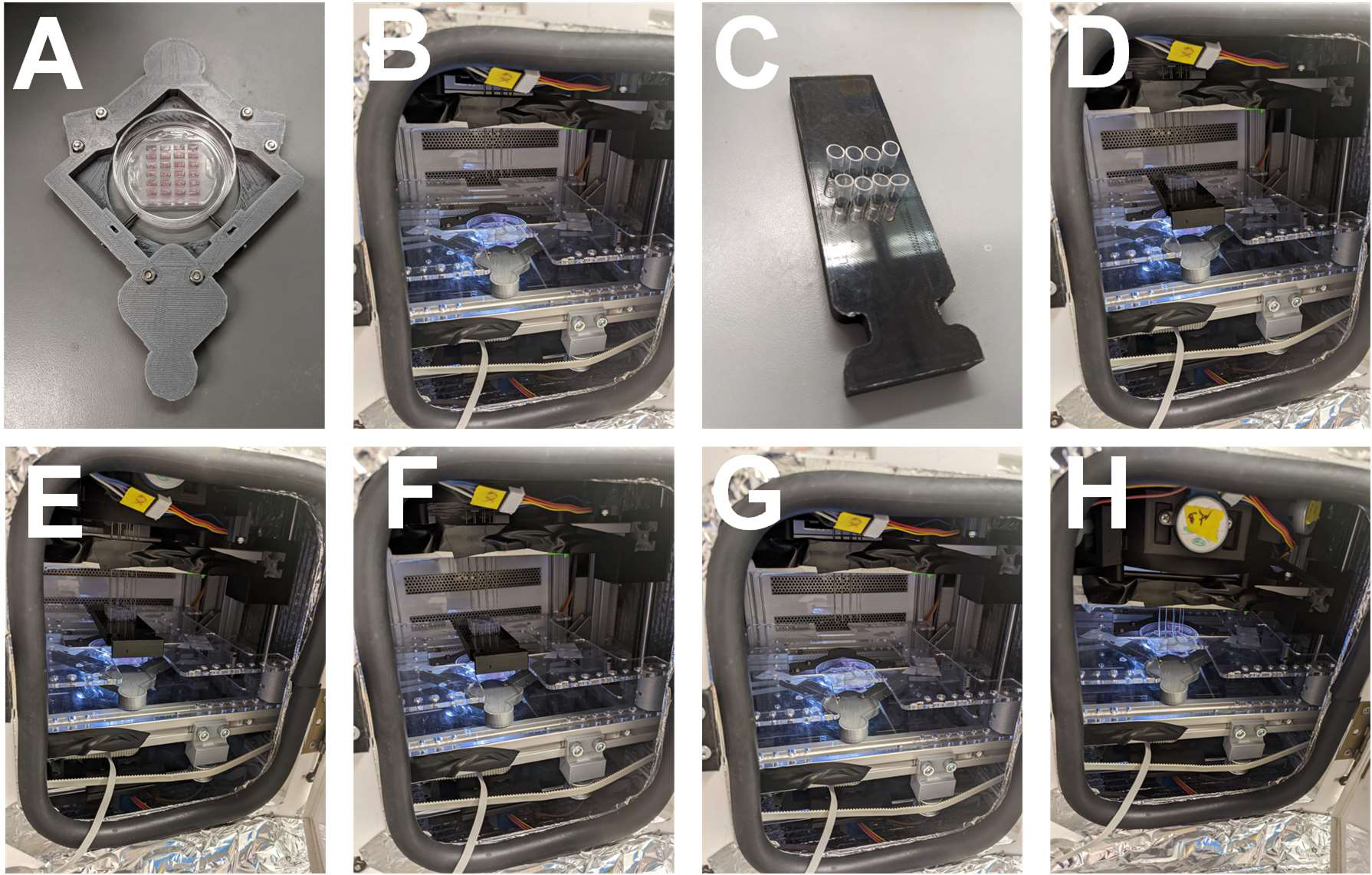
Seeding Process. **A)** Chip in chip holder. **B)** Holder inserted into the machine. **C)** PCR tubes in tube holder. **D)** PCR tubes filled with cell laden media and inserted into the machine. **E)** Cell dispensers resuspend cells and withdraw them. **F)** Cell Dispensers raised. **G)** Tube holder removed. **H)** Cell dispensers descend into the chip.

Flows generated by the syringe pump were modulated by the flexibility of the fluidic system, which depended on the size of the air gap between the priming water and the cell laden media, in addition to the flexibility of the Tygon tubing. Seeding parameters also depended on the clumping behavior of the cells during the seeding process. Larger clumps of cells are denser, and will sediment faster than smaller clumps or single cells. Large clumps of cells sometimes clog the dispensers, requiring additional flow to force them out, while small clumps may sediment under gravity without flow. Single cells take much longer to sediment, and require flow to enter the chip. iPSC passaging is performed using ReLeSR, a reagent that is designed to allow the cells to clump, which leads to higher viability. Cardiomyocytes and endothelial cells are removed from their plates using trypsin, resulting in smaller clumps or single cells. For cells that tend to clump, such as iPSCs, pulsatile flow was found to be more effective at moving them than continuous flow. iPSC clumps can easily clog dispensers, and resist being pushed into channels. They can be dislodged with bursts of flow of 0.5 μl at 9 μl/min, with one burst every 30 s. This is equivalent to an average flow of 1 μL/min, but a continuous flow does not dislodge clumps efficiently from the dispensers. While the syringe pump performed bursts at 9 μl/min, the movement was coupled to the cells via an air gap within the tubing and the compliance of the Tygon tubing material, resulting in a lower flow speed that extends over a longer length of time.

### Flexure Micromanipulators

Control over the X-Y position of each dispenser was necessary for controlling the seeding process. The cells were seeded into 800 μm diameter organoid culture wells in the microfluidic chip, and at minimum, each dispenser had to be accurately positioned in order to enter the holes. As the final cell dispenser diameter chosen was 762 μm (**Figure 7E**), each cell dispenser must be aligned to within 19 μm to enter the culture well at all, and additional adjustments within the well were helpful to reduce cell position biases that could impact organoid compaction later in development.

While a conventional X-Y gantry using belt or screw mechanisms with sliding linear bearings may have sufficed for a single cell dispenser, simultaneously seeding eight organoid culture wells in one chip requires eight densely packed gantries, and the height and complexity of conventional gantries would make this difficult. However, since the range of motion of each cell dispenser could be limited to a few millimeters, a flexure system was feasible.

Flexures are mechanisms that leverage the bending of material within a single part, rather than sliding of multiple parts. High precision flexure stages are commonly used in semiconductor fabrication instruments, atomic force microscopes, MEMS sensors and actuators, and other sub-micron control systems, as motion control below 100 nm is hard to control with conventional mechanisms.^32^ Flexures are also used at the other extreme of the cost spectrum, where they are used to make plastic hinges that can be injection molded as a single piece, reducing cost by eliminating assembly of multiple parts.^33^ Recently, 3D printing has been applied towards making complex flexure systems, such as the OpenFlexure microscope, in which a 3 axis flexure stage is printed as one piece using a consumer 3D printer.^30^ These mechanisms are limited in their range of motion, but when that limitation is acceptable, they have advantages in size, reliability, and precision.^34^

An X-Y flexure stage was designed based on a double parallelogram flexure element (**Figure 10A-B**), with the initial design adapted from Awtar et al.^32, 34^ Double parallelogram flexures act as linear bearings with long ranges of motion relative to other flexure systems;^34^ in our case, the micromanipulators have 6 mm ranges of motion. In order for the flexure micromanipulators to be cheap and easy to prototype in-house, they were 3D printed on an Ultimaker S3, with the CAD model and print slicing adjusted until the flexure elements were printed via a single pass from the print head. This resulted in flexure elements that are approximately 0.5 mm thick and 8 mm high. With an additional 2 mm spacing to ensure flexure stages would not rub against each other, the total gantry height was 10 mm, and the stack of 8 gantries controlling 8 micromanipulators was only 80 mm high. Unlike a conventional gantry, which requires time spent assembling and then adjusting all the components to ensure the gantry is aligned, level, and can move smoothly, the flexure gantries require minimal assembly and no alignment.

**Figure 10.**
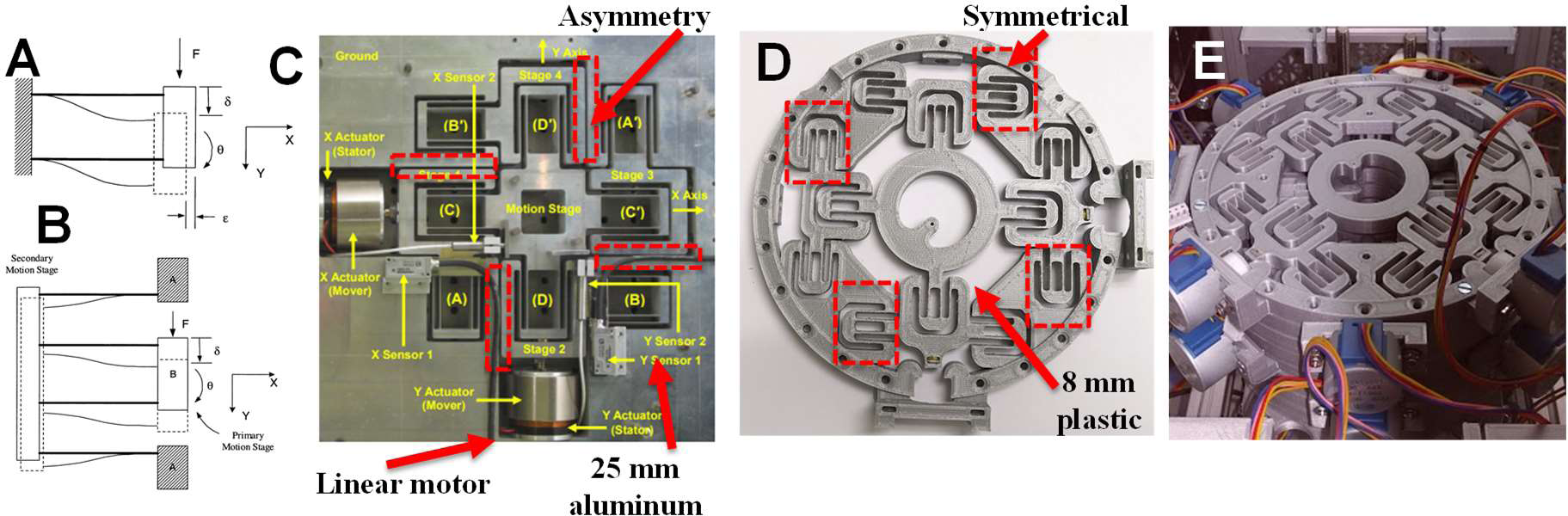
Flexure XY Micromanipulator Stage. **A)** Parallelogram flexure, with low resistance in the Y direction and high resistance in X. The stage remains parallel, but travels in an arc with off target motion in the –X direction. **B)** Double parallelogram flexure, with arcing motion by one set of flexures cancelled by the motion of the second set of flexures. Motion of B relative to A remains both parallel and linear. **C)** Asymmetrical aluminum flexure stage, with out-of-plane forces handled by the use of linear motors. **D)** Custom 3D printed plastic flexure stage, with out-of-plane forces handled by four additional flexures. **E)** A stack of 8 flexure gantries. **A**, **B**, and **C** adapted from Awtar et al.

The first version of the flexure stage was a direct copy of the configuration used by Awtar et al.,^32, 34^ with 8 flexures units per gantry (**Figure 10C**). This system has an asymmetry with linear bearings supported only on one side. As the Awtar micromanipulator system was designed to be used with linear voice coil motors that generate minimal out-of-plane forces, an asymmetrical bearing could be acceptable; in addition, the 25 mm height of their flexures made rejection of out of plane forces easier. However, for our low-cost adaptation of the flexure stage, motion was driven by a screw turned by a rotating stepper motor, generating considerable out-of-plane forces that impacted the 8 mm high 3D printed flexures. To increase the flexure stage’s ability to reject out-of-plane forces, a symmetrical flexure configuration was used with 12 flexures in total (**Figure 10D**). As the entire stage was 3D printed as a single piece, increases in geometric complexity did not increase cost or alignment complexity, though the added flexure units did increase the X-Y size of the flexure gantry from a 75 mm diameter to an 80 mm diameter. The complete stack of flexure gantries is shown in **Figure 10E**, and is shown with the cell dispensers and fluidics attached in **Figure 2B**. Connected to two vertical leadscrews, it could be raised to allow cells to be inserted into the machine, or lowered to withdraw the cells or deposit them into a chip (**Figure 9D-H**).

The flexure units were driven by 16 28BYJ-48 stepper motors with internal gearing resulting in 4096 steps per revolution. Ungeared stepper motors require power to hold their position, but gears add considerable friction to them, meaning that they can be powered down when not in use, and the positioned micromanipulator will maintain its position. The steppers turned M3 screws with 0.5 mm pitches, resulting in a final resolution of 0.122 μm/step, which is sufficient for fine positioning of the dispenser tips within the central organoid wells.

While repeatedly bending material back and forth could lead to material failure with large ranges of motion, the small range of motion bending used in flexure mechanisms leads to minimal material stress. The flexure systems used in this project were usable for at least 5 years without any material failures. However, if positioned in a stressed configuration for long periods of time, the plastic flexures would eventually relax into that position, which could lead to an increase in the force necessary to drive it to other positions. This may be exacerbated by th flexures being printed with thermoplastics and used at 37 °C. To remedy this problem, after every cell seeding experiment, the flexures are driven back to their center point to prevent warping.

Low-cost screw mechanisms require flexible shaft couplers to couple torque from the motors to the screws, as the motor and the screw can’t be expected to be aligned precisely. As no flexible shaft coupler was available that was appropriate for the size of this system, a custom 3D printed coupler was designed that uses hard ABS pieces coupling to the motor and the screw, with a soft TPU (Thermoplastic Polyurethane) coupler in between, providing the necessary flexibility (**Figure 11A-B**). For vertical adjustment of the cell dispensers, two TPU rubber washers were printed and glued to the XY stage, holding the dispensers at adjustable heights via friction (**Figure 11C**).

**Figure 11.**
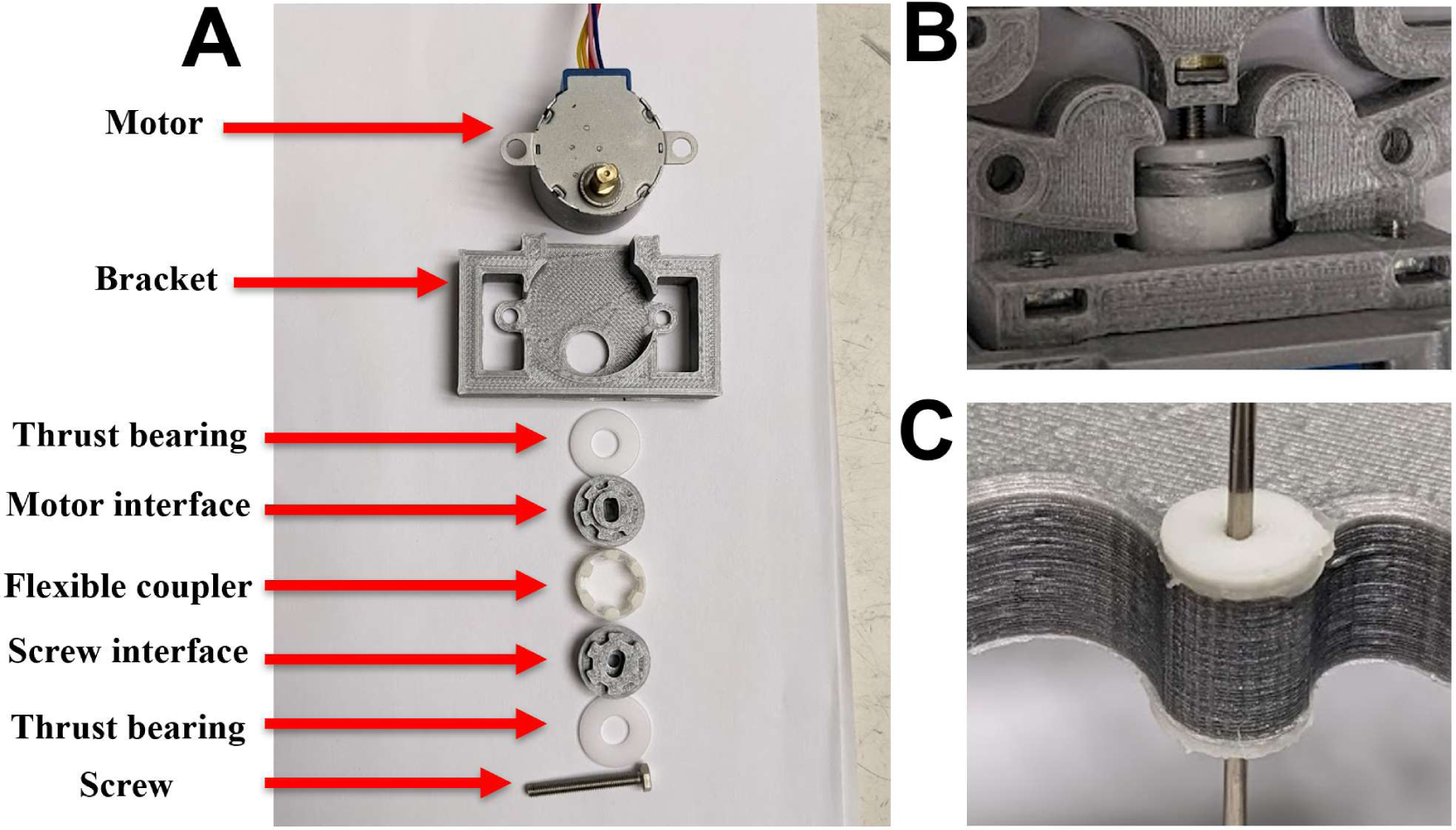
Motor Assembly for Flexure Micromanipulator Stage. **A)** Actuator components with 3D printed flexible shaft coupler. **B)** Assembled actuator system. **C)** TPU rubber washers allowing vertical adjustment.

### Time Lapse Imaging of Seeding Process

Due to machining tolerances, flexure warping over time, backlash of the motor gearing systems, and shifting of the dispensers, the micromanipulator positions were not very repeatable, despite being very high resolution. However, when controlled during visualization with the gantry microscope, dispensers could easily be positioned to their desired locations (**Figure 12A-B**). Time lapse imaging during a complete seeding is shown in **Figure 12**, with representative images from all 8 organoid culture wells. First, the cell dispensers were lowered into the chip, where they appeared above the organoid culture wells, but misaligned and out of focus due to being in a higher focal plane (**Figure 12A**). The micromanipulators were used with visual feedback to individually align each cell dispenser above its well (**Figure 12B**), with the exception of dispenser #8, which happened to already be aligned. The cell dispensers were lowered into the wells, where the seedings begin (**Figure 12C**). An intermediate point during the seeding is shown in **Figure 12D**, and **Figure 12E** shows the final cell distribution, with cells visible in every channel. This demonstrates the robotic seeding system’s ability to produce controlled cell seedings via independent X-Y control of each micromanipulator, vertical control of the entire micromanipulator block, environmental conditions, and syringe pump generated flow, all while observing via a microscope on a motorized gantry.

**Figure 12.**
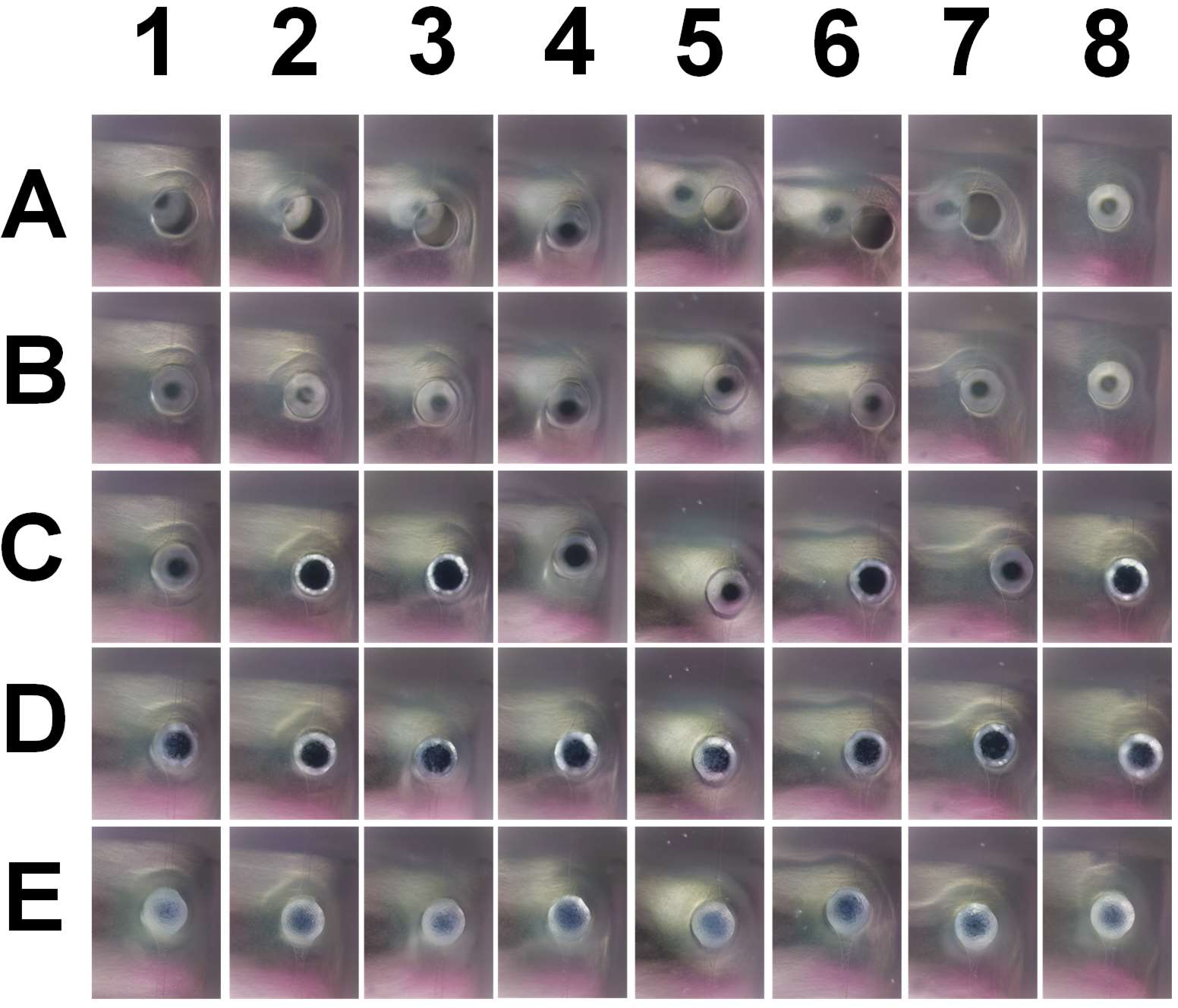
Time Lapse Imaging of Seeding Process. **A)** 8 cell dispensers are lowered into the chip, initially misaligned. **B)** Micromanipulators are used to independently align each cell dispenser. **C)** Cell dispensers are lowered into the chip, and the seeding process begins. **D)** The seeding process continues, and cells are observed to fall into the central organoid well, or are pushed into channels. **E)** The final configuration of cells when the seeding process is finished.

## Discussion

Before a microfluidic tube-shaped organoid can be grown in a chip, cells must be removed from a standard culture plate and deposited into the chip. While other microfluidic platforms have implemented this task successfully, and usually with simple techniques that were easily replicated, this simplicity is a function of the goal of the project, which is typically the production of spheres, surfaces lined with cells or patterns of cells, or the chaotic growth of angiogenic tubes. The task of growing larger, coherent, tube-shaped organoids revealed itself to be suprisingly subtle and difficult, and highly dependent on the intial positions of cells in the chip. Without controlled, repeatable seedings, it is unlikely that this venture will produce biological results with enough repeatability to lead to relevant conclusions.

As environmental control is known to be a critical factor in cell seeding, the cell seeding robot was embedded in a commercially available open hardware CO_2_ incubator, Incuvers, which was then modified to allow for on-the-fly user control over temperature, CO_2_ and humidity. The frame was rebuilt such that a larger robot could fit within it, it could be housed within a laminar flow hood to keep the process sterile, and a variety of access points were available for the inevitable maintenance of the internal machinery.

As this project demanded increasingly precise control over the seeding process, a microscope on a gantry was developed that used an oblique lighting system to monitor the entire seeding process, from the chip and cell dispenser alignments to the cells falling into the chip and recovering afterwards. As cells were observed entering the chip under different flow conditions produced by a syringe pump, with varying microfluidic chip architectures and cell dispenser dimensions, it became clear that the system would develop unexpected residual flow patterns that diverted cells away from the target. To address this problem, even more control over the system was added in the form of motor control over the vertical positions of the cell dispensers, as well as individual X-Y control over their positions with micromanipulators. An existing flexure gantry was modified, converting it from a thick, metal mechanism to a low cost 3D printed version that was stackable. These stacks of flexure micromanipulators allowed the cells to be guided to desired locations with greater accuracy and repeatability.

The completed robotic seeding system allowed for user or automated control over temperature, CO_2,_ humidity, flow, cell dispenser height, and the individual X-Y position of each cell dispenser, providing control over the seeding process.

